# Spatiotemporal dynamics of *Hemerocallis citrina* habitat suitability in China under climate change scenarios

**DOI:** 10.1101/2025.01.02.631172

**Authors:** Liu Yang, Yi Wu, Song Sheng, Jiqing Peng, Hanqing Qiu, Yuancheng Wen, Zhongyuan Ma, Jianjun Li, Sen Wang

## Abstract

**Objective:** *Hemerocailis citrina*, a perennial herb belonging to the *Hemerocailis* genus in the Apophaceae family, has been utilized as both food and medicine for thousands of years. Additionally, it is considered a traditional dominant cash crop in China. Therefore, analyzing the spatio-temporal distribution characteristics of climate sustability of *H. citrina* in current and future climatic conditions is crucial for its stable industry development in China.

**Method:** ENMTools was used to screen the data of 287 distribution points of *H. citrina*. 37 environmental variables affecting the distribution of *H. citrina* were selected, the MaxEnt model and Geographic Information System (ArcGIS) were used to simulate the potential distribution patterns of *H. citrina* in 5 periods (current, future 2040 s, 2060 s, 2080 s, and 2100 s), and the dominant climate factors affecting the distribution of *H. citrina* were analyzed through contribution rate, displacement importance value, and jackknife test.

**Results:** (1) The average AUC value of 10 cross-validated fits under the ROC curve for the training dataset was 0.933, and the average AUC value of 10 cross-validated fits for the test dataset was 0.901, this indicates that the simulation stability and reliability are highest when incorporating all three types of environmental variables. (2) The main environmental factors identified affecting their distribution include the lowest temperature of the coldest month, average temperature of the coldest quarter, slope, elevation, and carbonate or lime content, as determined by the contribution rate of cutting method, displacement contribution rate, and single factor response curve. (3) The model prediction showed that *H. citrina* was widely distributed in 34 provincial regions of China under the current climate conditions, with a suitable area of 646.81 km^2^. (4) Under the future emission scenarios, the suitable habitat for *H. citrina* is projected to shrink and not shift to higher latitudes, indicating a negative impact of climate warming on its distribution, which will reduce the suitable habitat and narrow the ecological niche. Temperature and precipitation have been identified as important factors affecting the distribution of *H. citrina*. Although *H. citrina*’s suitable habitat is expected to decrease to varying degrees in the future, the distribution area is relatively stable and there will still be a broad natural habitat in the future.

**Conclusion:** The species distribution model was used to predict the future temporal and spatial distribution of climate-suitable areas for *H. citrina* in China. Based on the model’s simulation results, it is recommended that north and southeast China become the main planting areas for *H. citrina* in the future, with Sichuan and other regions as key development areas. Additionally, other regions should prioritize developing suitable cash crops based on local conditions.

## 1 Introduction

The geographical distribution of species is shaped by the interplay of climate, soil, topographic, ecosystems, and human activities throughout their evolutionary history, reflecting their phylogenetic background, dispersal processes, and adaptability to new environments(Soberon et al.,2009). Since the 20th century, rapid population growth, coupled with accelerated industrialization and urbanization, has led to global warming and a decline in ecological quality, profoundly impacting species distribution patterns and the structure, function, and resilience of ecosystems(Gibson et al.,2011; Dieleman et al.,2015; Chen et al.,2011). Consequently, investigating the potential geographical distribution of species has emerged as a focal area in the fields of regional ecology and biogeography(Bellard et al.,2012). *H. citrina*, a perennial herb from the Apophaceae family, holds a longstanding cultural significance in China, serving both culinary and medicinal purposes for millennia. As a cornerstone of traditional agriculture, *H. citrina* not only symbolizes a primary cash crop but also plays a crucial role in China’s vegetable safety and industry. Thus, examining the temporal and spatial distribution of *H. citrin*a’s suitable habitats under present and future climatic scenarios is crucial for ensuring the sustainable progression of China’s *H. citrina* sector.

The species distribution model (SDM) integrates the geographic locations of target species populations with environmental variable data, employing specific algorithmic principles to estimate their ecological niches, subsequently projecting these niches into the environment. This process quantifies species’ habitat preferences as probabilistic measures(Zhuang et al.,2018). A diverse array of SDMs have been developed, leveraging various algorithmic principles. Prominent among these are the Climax(Araujo et al.,2018), GARP (Genetic Algorithm for Rule-set Production)(Elith et al.,2009), MaxEnt (Maximum Entropy)(Zhu et al.,2013), Bioclim(Elith et al.,2011), and Domain(Phillips et al.,2006) models. Integration of these models’ predictive outputs with ArcGIS software facilitates the generation of intuitive visual representations.

The MaxEnt model, predicated on the principle of maximum entropy, is particularly adept at extrapolating predictions from incomplete information. It determines the optimal probability distribution-representing the species’ most likely distribution under given constraints-by analyzing the non-random associations between species’ spatial distribution data and the climatic, topographic, and soil variables of the study area, thereby constructing a spatial distribution model on a geographical scale(Li et al.,2020). Advances in computer technology, remote sensing, ecology, and statistics have significantly enhanced the application scope and effectiveness of SDMs. The MaxEnt model, having undergone several refinements, is now extensively utilized in predicting the spatial distribution of species under current and future climatic conditions(Rivera-Parra et al.,2020).

Currently, China leads globally in the cultivation of *H. citrina* prompting numerous studies on its suitable growth areas within specific localities(Liu et al.,2020;Zhang et al.,2018). However, nationwide research on this subject remains limited, with scant attention given to how these suitable areas might shift under future climate scenarios. Moreover, integrated simulations employing multi-factorial species distribution models to assess *H. citrina*’s optimal growing regions are sparse. Utilizing Geographic Information System (GIS) and the MaxEnt model, this study leverages climate, soil, and topographic data from 287 *H. citrina* sample sites to systematically evaluate the potential suitable distribution areas for this species across China. It also identifies the key climatic factors influencing these areas and delineates the characteristics of regions with high suitability. This research is instrumental in comprehending the impact of climate change on the viability of *H. citrina* cultivation in China, thereby offering a theoretical framework to guide the strategic regionalization of *H. citrina* cultivation and to formulate responses to climate variability.

## 2 Materials and methods

### 2.1 Sources of data

#### 2.1.1 Geographic information of *H. citrina* population

The geographic data for the *H. citrina* population were sourced from the Global Biodiversity Information Facility (GBIF, [https://www.gbif.org/]), encompassing 1,383 specimen records for geographic analysis.

#### 2.1.2 Environmental variables information

This study considered three types of environmental variables:

(1) Climatic variables: Historical climate data (1970-2000) for 19 bioclimatic variables (Bio1*-*Bio19) were obtained from the WorldClim database ([https://www.worldclim.org/]). Future climate scenarios for 2030 (2021*-*2040) and 2050 (2041*-*2060) were also considered, based on the SSP245 scenario from the SSP*-*CSM2*-*MR climate model.
(2) Topographic variables: These include elevation, slope direction, and slope topography factors, sourced from WorldClim with a resolution of 2.5’.
(3) Soil variables: 15 soil factors were retrieved from the Harmonized World Soil Database (HWSD, [https://www.fao.org/soils-portal/en/]). Detailed information and coding of these variables are presented in Table 1.

**Table 1.** Environment variable.

| Code | Description | Code | Description |
| --- | --- | --- | --- |
| Bio<sub>1</sub> | Annual mean temperature | Ele | Elevation |
| Bio<sub>2</sub> | Mean diurnal range | Slo | Slope |
| Bio<sub>3</sub> | Isothermality | Asp | Aspect |
| Bio<sub>4</sub> | Temperature seasonality | SP<sub>1</sub> | Percentage of gravel volume |
| Bio<sub>5</sub> | Maximum temperature of the warmest month | SP<sub>2</sub> | Sand content |
| Bio<sub>6</sub> | Minimum temperature of the coldest month | SP<sub>3</sub> | Silt content |
| Bio<sub>7</sub> | Temperature annual range | SP<sub>4</sub> | Clay content |
| Bio<sub>8</sub> | Mean temperature of the wettest quarter | SP<sub>5</sub> | Soil bulk density |
| Bio<sub>9</sub> | Mean temperature of the driest quarter | SP<sub>6</sub> | Organic carbon content |
| Bio<sub>10</sub> | Mean temperature of the warmest quarter | SP<sub>7</sub> | pH value |
| Bio<sub>11</sub> | Mean temperature of the coldest quarter | SP<sub>8</sub> | Cohesive layer soil cation exchange capacity |
| Bio<sub>12</sub> | Annual precipitation | SP<sub>9</sub> | Soil cation exchange capacity |
| Bio<sub>13</sub> | Precipitation of wettest month | SP<sub>10</sub> | Basic saturation |
| Bio<sub>14</sub> | Precipitation of driest month | SP<sub>11</sub> | Exchangeable bases |
| Bio<sub>15</sub> | Precipitation seasonality | SP<sub>12</sub> | Carbonate or lime content |
| Bio<sub>16</sub> | Precipitation of the wettest quarter | SP<sub>13</sub> | Sulfate content |
| Bio<sub>17</sub> | Precipitation of the driest quarter | SP<sub>14</sub> | Exchangeable sodium salt |
| Bio<sub>18</sub> | Precipitation of the warmest quarter | SP<sub>15</sub> | Conductivity |
| Bio<sub>19</sub> | Precipitation of the coldest quarter |  |  |

### 2.2 Data processing

From the 1 383 geographic specimen records, duplicates, incorrect coordinates, and records lacking detailed geographic information were removed. Using ArcGIS 10.8, data points outside China were excluded, and proximity analysis was conducted to average out points within a 10 km^2^ area, ENMTools was used to screen the data of 287 effective distribution points of *H. citrina*. These were saved as a .CSV file in Excel 2016. The environmental variables were standardized to the WGS1984 geographic coordinate system in ArcGIS, with dimensions of 1 646 by 1 220 rows and columns, respectively. Using the vector map No. GS(2019)1822 of the standard map of China for reference, the environmental factors specific to China were extracted using the masking tool in ArcGIS and converted into ASCII format for MaxEnt software analysis. This process facilitated the extraction of environmental variable data for the 287 *H. citrina* samples, with key variables retained for the final modeling in the MaxEnt model.

### 2.3 Selection and reliability test of model parameters

For setting the parameters of the MaxEnt model, we followed the methodology of Wang(Wang et al.,2023), allocating 75% of the data for the training set and limiting the maximum number of iterations to 10,000. The Bootstrap method was employed for resampling, and options like Jackknife and Response Curves were selected to generate response curves, enhancing the model’s analytical depth. To minimize the impact of outliers, the number of replicates was set at 10, and the average of 10 iterations was calculated to ensure stable simulation outcomes(Rivera-Parra et al.,2020). The model’s accuracy was assessed using the Receiver Operating Characteristic (ROC) curve. The Area Under the ROC Curve (AUC) ratings-spanning from 0.5 to 1-categorize the model’s simulation performance as “poor” (0.5*-* 0.6), “fair” (0.6*-*0.7), “average” (0.7*-*0.8), “good” (0.8*-*0.9), and “excellent” (0.9*-*1)(Liu et al.,2020). In MaxEnt model predictions, the percentage of contribution is utilized to determine the significance of individual environmental factors, with higher values indicating greater impact on the model’s results. Permutation ordering assesses the influence of environmental variables based on a random assortment from the training sites and environmental data, where higher values signify a more substantial collection of environmental variables by the site. Among the output formats-Raw, Logistic, and Cumulative-the complementary log*-*log (Cloglog) model, based on the inhomogeneous Poisson process (IPP), was identified as having superior theoretical justification. Currently, the “Cloglog” output is considered the most effective for predicting the optimal suitability area(Zhang et al.,2018), offering a robust framework for environmental modeling in MaxEnt.

### 2.4 Environmental Importance assessment

To mitigate multicollinearity among environmental factors and prevent overfitting in the MaxEnt model, the ENMTool software was utilized for correlation analysis. A Pearson correlation coefficient threshold of 0.8 was established to identify and eliminate highly correlated environmental factors. In cases where two environmental variables exhibited a correlation coefficient of ≥0.8, the one with a higher contribution rate, as indicated by the initial MaxEnt model results, was retained [18]. Consequently, 22 environmental factors were selected for the MaxEnt analysis, encompassing various aspects of the maximum entropy model. These factors include Aspect (Slope), Slope (slope aspect), Elev (altitude), Bio2 (mean diurnal temperature range), Bio3 (isothermality), Bio4 (temperature seasonality), Bio5 (max temperature of warmest month), Bio6 (min temperature of coldest month), Bio12 (annual precipitation), Bio15 (precipitation seasonality), T_caso4 (sulfate content), T_gravel (gravel volume percentage), T_cec_clay (cation exchange capacity of clay soil), T_oc (organic carbon content), T_pH_H_2_O (soil pH), T_clay (clay content), T_bs (base saturation), T_ref_bulk (soil bulk density), T_caco3 (carbonate or lime content), T_esp (exchangeable sodium percentage), T_silt (silt content), and T_teb (total exchangeable bases). The model simulations were conducted using singular climate variable (Climate, C), combined climate and topographical variable (Climate and Topographical, C+T), and an integrated set of climate, topographical, hydrological, and soil variables (C+T+S). This facilitated a comparative analysis of the model’s response to different environmental variables, helping to discern the distinctive effects of each variable set on the simulation outcomes.

### 2.5 Division of suitable zones

In processing the MaxEnt simulation data, ArcGIS 10.8 software was employed to integrate the average of ten simulation runs and convert these into raster format. This raster data represents the survival probability of *H. citrina*, with each raster value indicating the likelihood of the species thriving in a given location. To classify the suitability zones, the balance training omission rate, predicted area, and threshold value (TPT) were used as criteria. Areas with a probability less than 0.1 were designated as non*-*suitable zones. A probability range greater than 0.1 but less than 0.3 was categorized as low suitability, greater than 0.3 but less than 0.6 as medium suitability, and greater than 0.6 as high suitability zones. The MaxEnt model’s output, specifically the distribution probability layer of *H. citrina*, was segmented into non*-*suitable, low suitable, medium suitable, and high suitable areas. These zones were quantified based on the actual pixel area to determine the extent of each suitability zone accurately.

## 3 Results

### 3.1 Model accuracy and stability test

Utilizing geographic data from 287 effective populations of *H. citrina* across China, the MaxEnt model simulations were conducted with three sets of comprehensive environmental variables: single climate variable (C), climate and topographic variables (C+T), and climate, topographic, and soil variables (C+T+S). The results, illustrated in Figure 1 and Table 2, demonstrate that the Area Under the Curve (AUC) values for the training sets are 0.907, 0.920, and 0.933 for C, C+T, and C+T+S, respectively, while the AUC values for the test sets are 0.895, 0.899, and 0.901, respectively. These figures indicate that the model incorporating all three types of environmental variables (C+T+S) offers the best simulation stability and high reliability, with a “preferably” predictive performance. Consequently, the MaxEnt model, when employing these three environmental variables, is shown to have high stability in simulating the potential distribution area of the *H. citrina*.

**Table 2.** Area under the receiver operating characteristic curve(AUC) under 3 types of environment variables.

| Environment variable type | Training AUC | Test AUC |
| --- | --- | --- |
| C | 0.907±0.001 | 0.895±0.012 |
| C+T | 0.920±0.001 | 0.899±0.012 |
| C+T+S | 0.933±0.001 | 0.901±0.125 |

topography and soil. The same below.

**Fig. 1.**
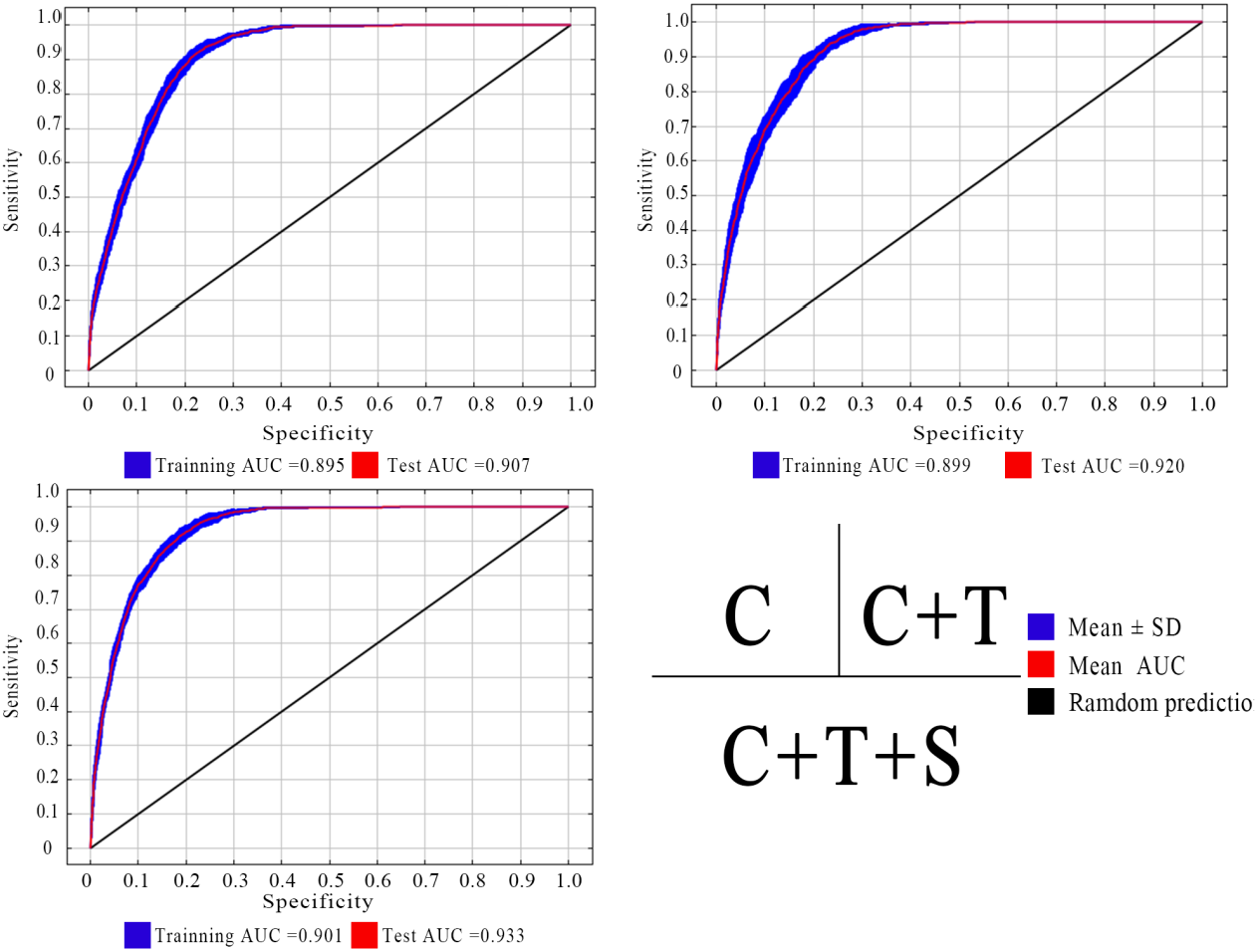
Model parameters of the model under 3 types of environment variables

### 3.2 Dominant environmental variables influencing the distribution of H. citrina

The analysis of *H. citrina* distribution incorporated three categories of environmental variables: single climate variable (C), Class 2 environmental variables (C+T), and Class 3 environmental variables (C+T+S). The Area Under the Curve (AUC) values for these variables, both for training and testing, are detailed in Table 2. Additionally, the contribution rates and importance values of these variables are presented in Table 3.

**Table 3.** Percent contribution of each environmental variable.

| Code | C+T+S |  | C+T |  | C |  |
| --- | --- | --- | --- | --- | --- | --- |
|  | Percent | Permutation | Percent | Permutation | Percent | Permutation |
|  | contribution/ | importance/% | contribution/ | importance/% | contribution/ | importance/% |
|  | % |  | % |  | % |  |
| Aspect | 2.0 | 2.2 | 2.5 | 1.9 |  |  |
| Elevation | 2.3 | 7.5 | 2.2 | 5.0 |  |  |
| Slope | 5.9 | 5.0 | 5.9 | 9.3 |  |  |
| SP<sub>3</sub> | 0.7 | 0.9 |  |  |  |  |
| SP<sub>13</sub> | 0.2 | 0.3 |  |  |  |  |
| SP<sub>7</sub> | 0.5 | 1.0 |  |  |  |  |
| SP<sub>12</sub> | 2.2 | 2.9 |  |  |  |  |
| SP<sub>1</sub> | 1.4 | 1.6 |  |  |  |  |
| SP<sub>11</sub> | 0.6 | 3.0 |  |  |  |  |
| SP<sub>5</sub> | 1.1 | 0.3 |  |  |  |  |
| SP<sub>9</sub> | 0.1 | 0.2 |  |  |  |  |
| SP<sub>8</sub> | 0.4 | 0.8 |  |  |  |  |
| SP<sub>10</sub> | 1.3 | 0.9 |  |  |  |  |
| SP<sub>11</sub> | 0.3 | 0.9 |  |  |  |  |
| SP<sub>4</sub> | 0.6 | 1.2 |  |  |  |  |
| SP<sub>6</sub> | 0.3 | 0.6 |  |  |  |  |
| Bio<sub>2</sub> | 0.8 | 0.8 | 3.0 | 2.5 | 4.3 | 3.4 |
| Bio<sub>3</sub> | 0.5 | 2.0 | 3.2 | 2.0 | 5.4 | 4.1 |
| Bio<sub>4</sub> | 0.3 | 0.2 | 1.9 | 10.5 | 2.0 | 16.1 |
| Bio<sub>5</sub> | 0.3 | 2.6 | 3.3 | 14.6 | 5.0 | 24.7 |
| Bio<sub>6</sub> | 0.2 | 2.0 | 35.4 | 27.1 | 31.3 | 10.1 |
| Bio<sub>12</sub> | 0.1 | 1.0 | 37.5 | 22.1 | 46.9 | 34.9 |
| Bio<sub>15</sub> | 0 | 0 | 5.0 | 5.1 | 5.2 | 6.6 |

From Table 3, it’s evident that under the comprehensive environmental scenario (C+T+S), the four most influential environmental variables based on contribution rate were: the lowest temperature of Annual mean water amount(Bio12) at 36.3%,the coldest month (Bio6) at 28.3%, Slope at 5.9%,Monthly mean of diurnal temperature difference (Bio2) at 4.5% Similarly, the variables with significant importance values were Annual mean water amount(Bio12) at 22.3%,the coldest month (Bio6) at 22.2%, Highest temperature of hotest month (Bio5) at 12.3%, Altitude (elev) at 7.5%.

Figure 2 displays the importance testing results of these variables using the AUC knife*-*edge method. In scenarios where only single variables were considered, the most critical environmental variables were identified as Bio12(Annual mean water amount), Bio6 (lowest temperature in the coldest month),,Bio2 (monthly mean diurnal temperature difference), Bio15(Variance of precipitation change),. Upon a comprehensive analysis of environmental variables’ contribution rates, replacement importance values, and the AUC knife*-*edge method, it was deduced that temperature (specifically during the annual mean water amount and lowest temperature in the coldest month), topographic (including slope and Elev), and soil factors (like carbonate or lime content and percentage of gravel volume) are the principal environmental factors constraining the distribution of *H. citrina*. This comparative analysis underscores that the MaxEnt model, using all three categories of comprehensive environmental variables (C+T+S), is more effective in identifying crucial environmental factors affecting *H. citrina* distribution, yielding more reliable simulation outcomes.

**Fig. 2.**
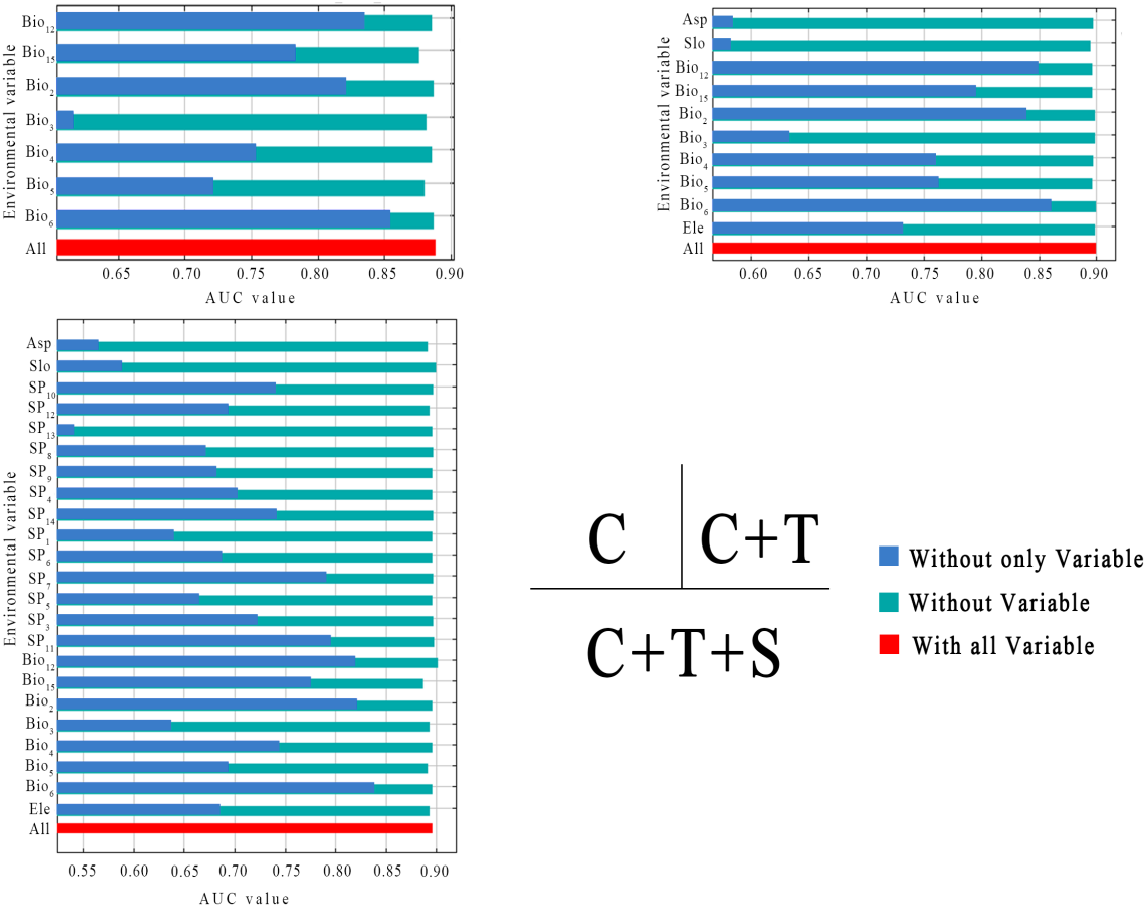
AUC values under the ROC curve of 3 types of environment variables

### 3.3 Potential suitable distribution area of H. citrina

Using ArcGIS 10.8 software, the simulation results for *H. citrina* were reclassified into four suitability categories: non*-*suitable areas (>0.1), low suitable areas (0.1–0.3), medium suitable areas (0.3– 0.6), and high suitable areas (>0.6). This classification facilitated a clearer visualization of the distribution patterns. The analysis incorporated single climate variable (C), combined climate and topographic variables (C+T), and integrated climate, topographic, and soil variables (C+T+S) to simulate *H. citrina*’s suitable areas, detailed in Table 4. The respective simulated suitable areas by these three environmental variable categories were 279.67 km^2^ (C+T+S), 300.93 km^2^ (C+T), and 319.67 km^2^ (C), indicating a decreasing trend in area from single to complex variable simulations.

**Table 4.**
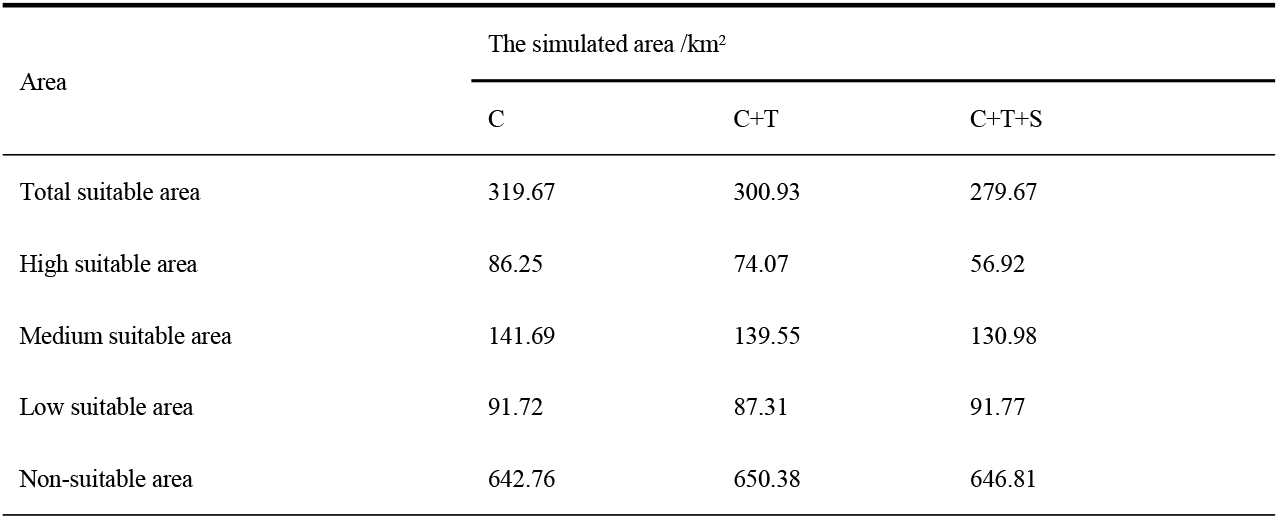
The suitable distribution area of *H. citrina*.

| Area | The simulated area /km<sup>2</sup> |  |  |
| --- | --- | --- | --- |
|  | C | C+T | C+T+S |
| Total suitable area | 319.67 | 300.93 | 279.67 |
| High suitable area | 86.25 | 74.07 | 56.92 |
| Medium suitable area | 141.69 | 139.55 | 130.98 |
| Low suitable area | 91.72 | 87.31 | 91.77 |
| Non-suitable area | 642.76 | 650.38 | 646.81 |

**Table 5.**
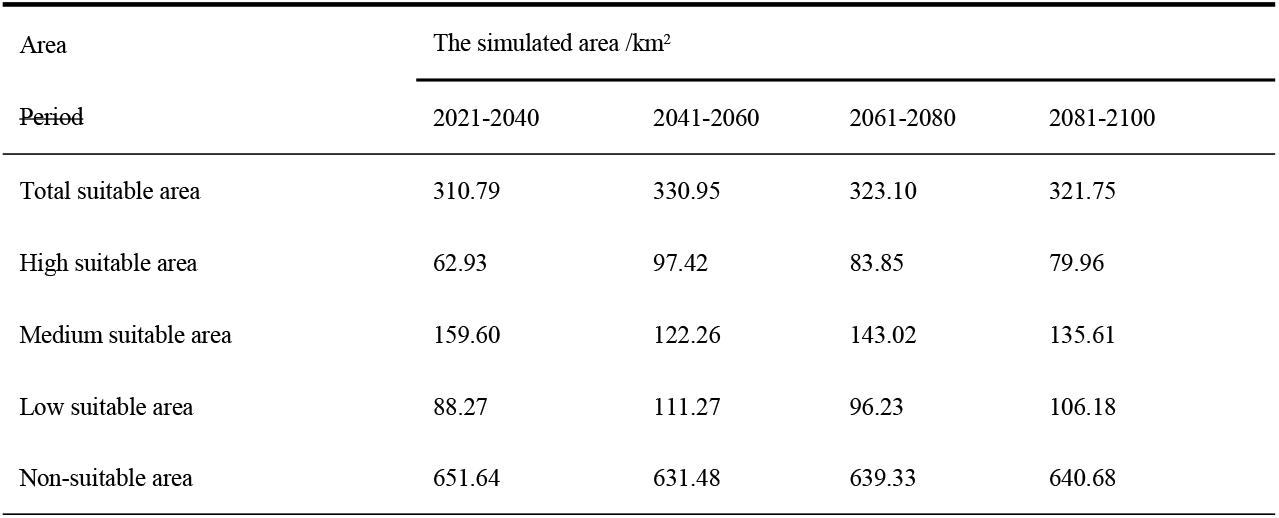
The suitable distribution area of *H. citrina* in future period.

| Area | The simulated area /km<sup>2</sup> |  |  |  |
| --- | --- | --- | --- | --- |
| Period | 2021-2040 | 2041-2060 | 2061-2080 | 2081-2100 |
| Total suitable area | 310.79 | 330.95 | 323.10 | 321.75 |
| High suitable area | 62.93 | 97.42 | 83.85 | 79.96 |
| Medium suitable area | 159.60 | 122.26 | 143.02 | 135.61 |
| Low suitable area | 88.27 | 111.27 | 96.23 | 106.18 |
| Non-suitable area | 651.64 | 631.48 | 639.33 | 640.68 |

The spatial projection in ArcGIS 10.8 was set to the Albers equal area conic projection, with the detailed outcomes illustrated in Figure 3. The simulations underscored the significant influence of climate factors on *H. citrina*’s geographic distribution: 100.1% contribution in single climate variable simulations (C), 89.3% in climate and topographic variable simulations (C+T), and 80.1% in combined climate, topographic, and soil simulations (C+T+S), underlining the paramountcy of climate factors in determining *H. citrina*’s growth.

**Fig. 3.**
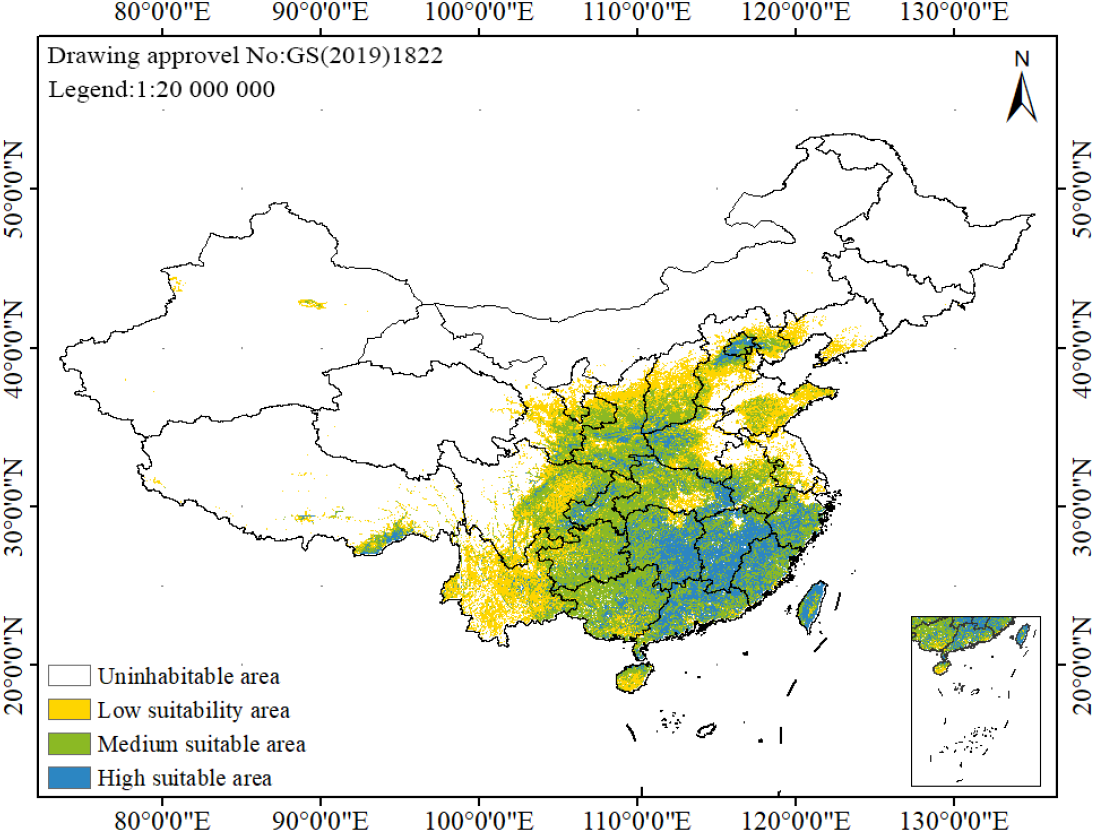
Suitable area distribution map of *H. citrina*

MaxEnt model predictions highlight that *H. citrina*’s suitable habitats predominantly span the eastern and southern regions of Hunan, as well as areas in Jiangxi, eastern Hubei, northwestern Henan, southern Shanxi, northern Shaanxi, and southwestern Gansu, totaling 279.67 km^2^. Distinctly, the most favorable habitats for Xanthoxanthium (apparently another species under discussion) were located in specific regions of Hunan, Hubei, Jiangxi, Henan, Shanxi, Shaanxi, Gansu, and central Anhui, with a notable suitable area of 56.92 km^2^, making up 20.35% of the total viable growth space. Additionally, substantial suitable habitats were identified in Guizhou, Guangxi, Yunnan, Hebei, and Anhui, totaling 130.98 km^2^ or 46.83% of the overall suitable area. Conversely, low suitability zones were primarily in Hainan, Jilin, Heilongjiang, accounting for 91.77 km^2^ or 32.81% of the assessed area. Non*-*suitable zones included Tibet Autonomous Region, Qinghai, and Xinjiang Uyghur Autonomous Region, covering 646.81 km^2^.

### 3.4 Simulation of suitable growth area of H. citrina under future climate scenarios

The simulation of *H. citrina* suitable growth areas under future climate scenarios reveals significant changes compared to historical data (Fig. 4). For the period 2021-2040, there is a notable increase in both the total and highly suitable areas compared to the historical period (1997-2000), with the total suitable area expanding from 279.36km^2^ to 310.79 km^2^. The high suitability area also saw an increase from 56.92 km^2^ to 62.93 km^2^, nearly doubling in size.

**Fig. 4.**
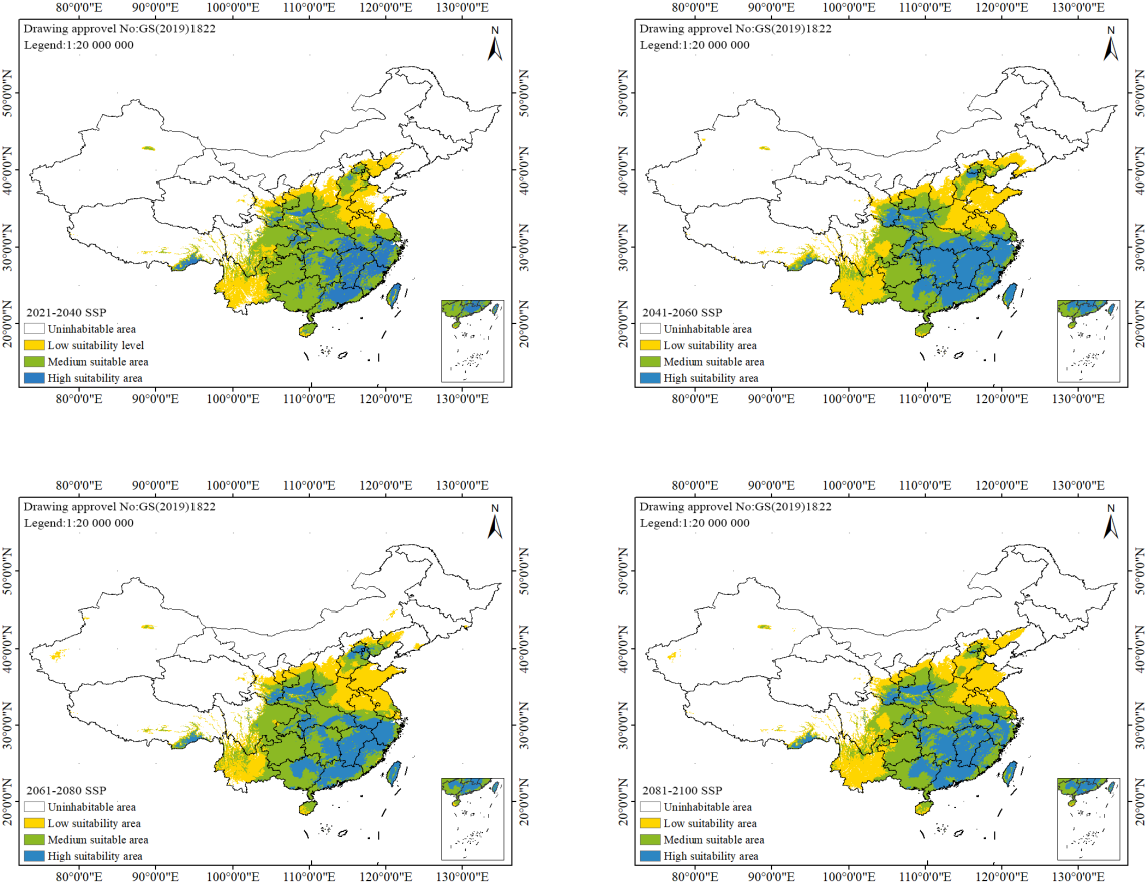
Suitable distribution area of *H. citrina* in future period

**Fig. 5.**
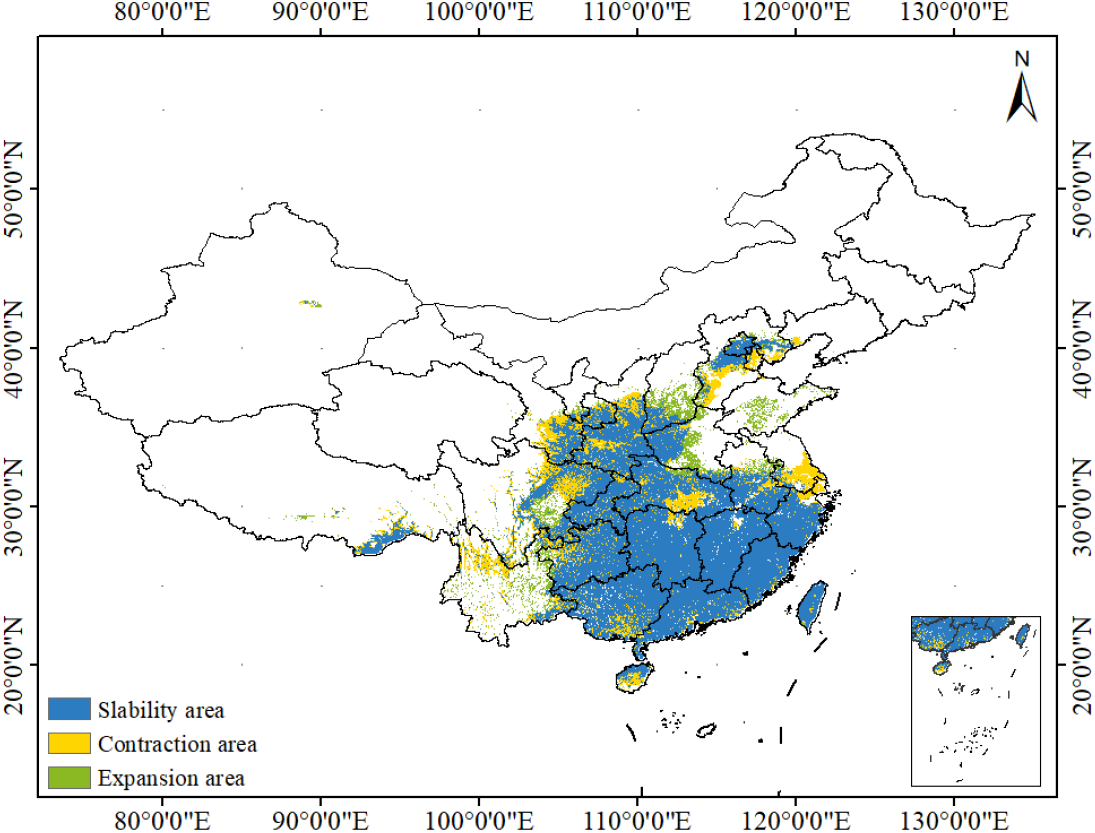
Distribution of stable expansion and contraction zones of *H. citrina*

Looking ahead to 2041-2060, the total suitable area further expands to 330.95 km^2^, significantly exceeding the earlier period (2021-2040), while the non-suitable area diminishes, contracting from 651.64 km^2^ to 631.48 km^2^. This trend suggests a future reduction in non-suitable areas and an expansion of suitable regions for *H. citrina* cultivation. According to the Intergovernmental Panel on Climate Change (IPCC) fifth report, global average surface temperatures are projected to rise by 0.3 to 4.8°C by the end of the 21st century, potentially altering species’ geographical distribution patterns[6].

The analysis indicates dynamic shifts in *H. citrina*’s suitable areas across different future periods, reflecting the potential impact of varying climate conditions. In the SSP245 scenario, the centroid of the suitable area is projected to move northwest by 732 km in the 2040s from its current position. In the 2060s, it shifts northwest by 108 km, followed by a northwest migration of 106 km in the 2080s. In the 2100s, the centroid moves northeast by 208 km, signifying a gradual transition from Hubei to Sichuang province over the course of the century. These migrations highlight the shifting dynamics of *H. citrina*’s suitable growing regions in response to future climate scenarios.

## 4 Discussion

### 4.1 Reliability of model simulation results

The MaxEnt model, leveraging maximum entropy theory, excels in simulating the relationship between species distribution and environmental factors. Notably, it demonstrates superior performance in predicting distributions from small sample sizes, where MaxEnt’s advantages become particularly evident(Elith et al.,2006;Chen et al.,2021;Wang et al.,2020). In this research, 287 samples were utilized, effectively mitigating the overfitting issue common in small sample analyses, thereby ensuring the accuracy and stability of the simulations(Zhao et al.,2018;Phillips et al.,2006). Running 10 iterations with the MaxEnt software yielded AUC values for both the training and test datasets close to or reaching 0.9, indicating high reliability and precision.

Geographical analysis revealed that *H. citrina* predominantly thrives in central and southern China, aligning with the “Flora of China”, which notes *H. citrina*’s distribution in provinces south of the Qinling Mountains, including southern Gansu, Shaanxi, and extending to Hebei, Shanxi, and Shandong(Editorial Committee of Flora of China, 1978). Furthermore, research from the Beijing Botanical Garden confirms that *H. citrina* mainly occupies areas south of 60°N, consistent with the simulation results of this study(Wang et al.,2014). The Guangdong Academy of Agricultural Sciences, considering both cultivation and market conditions, identifies the cultivation advantages of the Chinese *H. citrina* industry in four regions: Northeast China, North China, the Yangtze River basin, and the Pearl River basin, pinpointing specific locations like Hunan (Qidong, Shaodong), Henan (Huaiyang), Shaanxi (Dali), Gansu (Qingyang), Shanxi (Datong), and Jiangsu (Suqian) as prime cultivation sites(Li et al.,2018;Fan et al.,2019;Fu et al.,2020). These findings closely match the high suitability areas for *H. citrina* identified in this study. Additionally, regions in Guangdong (Haifeng and Dapu), Yunnan (Xianguan), Fujian (Dehua), Zhejiang (Jinyun), Jiangxi, Guangxi, and Hainan are recognized for their high suitability, underscoring their potential for further development in *H. citrina* cultivation.

### 4.2 Main environmental variables influencing the distribution of H. citrina

The distribution of plant populations reflects the cumulative effects of biological and abiotic factors, embodying the plant’s adaptive response to the environment. The simulation employing comprehensive environmental variables (C+T+S) highlighted that air temperature (specifically the lowest temperature in the coldest month and seasonal temperature variation), topographic (including slope direction and elevation), and soil characteristics (like soil bulk density and pH value) are the critical environmental factors constraining the distribution of *H. citrina*. This finding aligns with *H. citrina*’s ecological adaptability to drought and infertile soils, with temperature playing a pivotal role. *H. citrina* thrives in warm conditions and exhibits robust adaptability; its aerial parts are frost-sensitive, but its subterranean components can overwinter in cold regions. The plant flourishes in temperatures between 14-20°C, especially during budding and flowering phases, where high temperatures and significant diurnal temperature variations promote vigorous growth and bud formation. Hydrothermal conditions largely dictate the distribution of plant species or vegetation types on a broad regional to global scale. However, at the landscape and smaller scales, local environmental factors predominantly influence vegetation distribution patterns(Yao et al.,2023). The spatial distribution of plant populations is influenced by a nexus of environmental factors, including temperature, moisture, and light, particularly across altitude gradients(Zhang et al.,2023). Tang’s research(Tang et al.,2020) indicates that elevation critically determines the distribution of mountain vegetation at the community scale across different geoclimatological zones. It is conjectured that topographic factors, primarily slope direction, along with precipitation, temperature, and soil conditions, integratively influence *H. citrina*’s geographical distribution pattern.

### 4.3 Distribution dynamics of H. citrina in response to climate change

Under the SSP245 climate change scenario, *H. citrina* exhibits a low rate of both expansion and contraction, with a consistent increase in its stable area. In comparison to the current suitable distribution, *H. citrina*’s suitable area has expanded, predominantly in the central, eastern, and southern China, stabilizing at 168.37 km^2^. Future climate projections suggest this area will maintain its suitability for *H. citrina*. The expansion and contraction zones are relatively modest in size. Expansion is primarily observed in the western and southeastern parts of Inner Mongolia, western Jilin, and the central and western regions of the Xinjiang Uyghur Autonomous Region, amounting to a minor growth of 42.60 km^2^. Conversely, contraction zones are located in southeastern Jilin, the central and southern areas of Inner Mongolia, and the northwestern part of the Xinjiang Uyghur Autonomous Region, with a total reduction area of 18.97 km^2^. This analysis underscores the nuanced impact of climate change on *H. citrina*’s distribution, with a notable trend towards stability and moderate shifts in its geographical range, reflecting the plant’s adaptability to evolving climate conditions.

## 5 Conclusions

(1) This study employed the MaxEnt model and ArcGIS to simulate and predict the distribution of *H. citrina* across China, categorizing the suitable areas into four levels: non-suitable, low suitable, medium suitable, and high suitable. The AUC value of the model reached 0.800, showcasing its relatively high accuracy.
(2) Additionally, the distribution of *H. citrina* was analyzed using the MaxEnt model and ArcGIS, revealing a wide distribution across north, central, and south China, as well as parts of northwest China, primarily east of the Mohe-Tengchong line. The Hoh Xil uninhabited area in Qinghai province and certain regions in Tibet and Xinjiang were identified as unsuitable for *H. citrina* cultivation.The lowest temperature of the coldest month is the most important influencing factor. *H. citrina*’s suitable area has expanded, predominantly in the central and southern China under climate change.
(3) The MaxEnt model, while focusing on climate, topographic, and soil as primary influencing factors, indicated that these elements significantly affect the growth and distribution of plants. However, there was a notable discrepancy between the climate-only simulated distribution of *H. citrina* and its actual distribution. Integrated environmental variables (climate, topographic, and soil) yielded simulations that more accurately represented *H. citrina*’s actual population distribution. Nonetheless, the determinants of plant distribution extend beyond these three factors. Light conditions, soil texture, species interactions, and human activity also play crucial roles in plant distribution. This study acknowledges its limitations due to the narrow focus on specific factors. Future research should incorporate a broader range of influences and examine more detailed scales to enhance the accuracy of the findings.

## Declaration of Competing Interest

The authors declare that they have no known competing financial interests or personal relationships that could have appeared to influence the work reported in this paper.

## Data availability

Data will be made available on request.

## Acknowledgments

This research was jointly supported by the National Key Research and Development Program(2022YFD2200505), Graduate Innovation Fund Project of Central South University of Forestry & Technology (2024CX02033).

## Notes

### Competing Interest Statement

The authors have declared no competing interest.

